# Oil Red O based method for Exosome Labelling and detection

**DOI:** 10.1101/2022.01.09.475514

**Authors:** Shikha Bharati, Km Anjaly, Shivani Thoidingjam, Ashu Bhan Tiku

## Abstract

With the realization of the role of exosomes in diseases especially cancer, exosome research is gaining popularity in biomedical sciences. To understand exosome biology, their labelling and tracking studies are important. New and improved methods of exosome labelling for detection and tracking of exosomes need to be developed to harness their therapeutic and diagnostic potential. In this paper, we report a novel, simple and effective method of labelling and detecting exosomes using Oil red O (ORO) which is a dye commonly used for lipid staining. Using ORO is a cost effective and easy approach with intense red colouration of stained exosomes. Further, the issues faced with commonly used lipophilic dyes for exosomes labelling such as long term persistence of dyes, aggregation and micelle formation of dyes, difficulty to distinguish dye particles from labelled exosomes and detection of large aggregates of dye or dye-exosome are not seen with ORO dye. This method shows good labelling efficiency of exosomes with very sensitive detection and real-time tracking of the cellular uptake of exosomes.

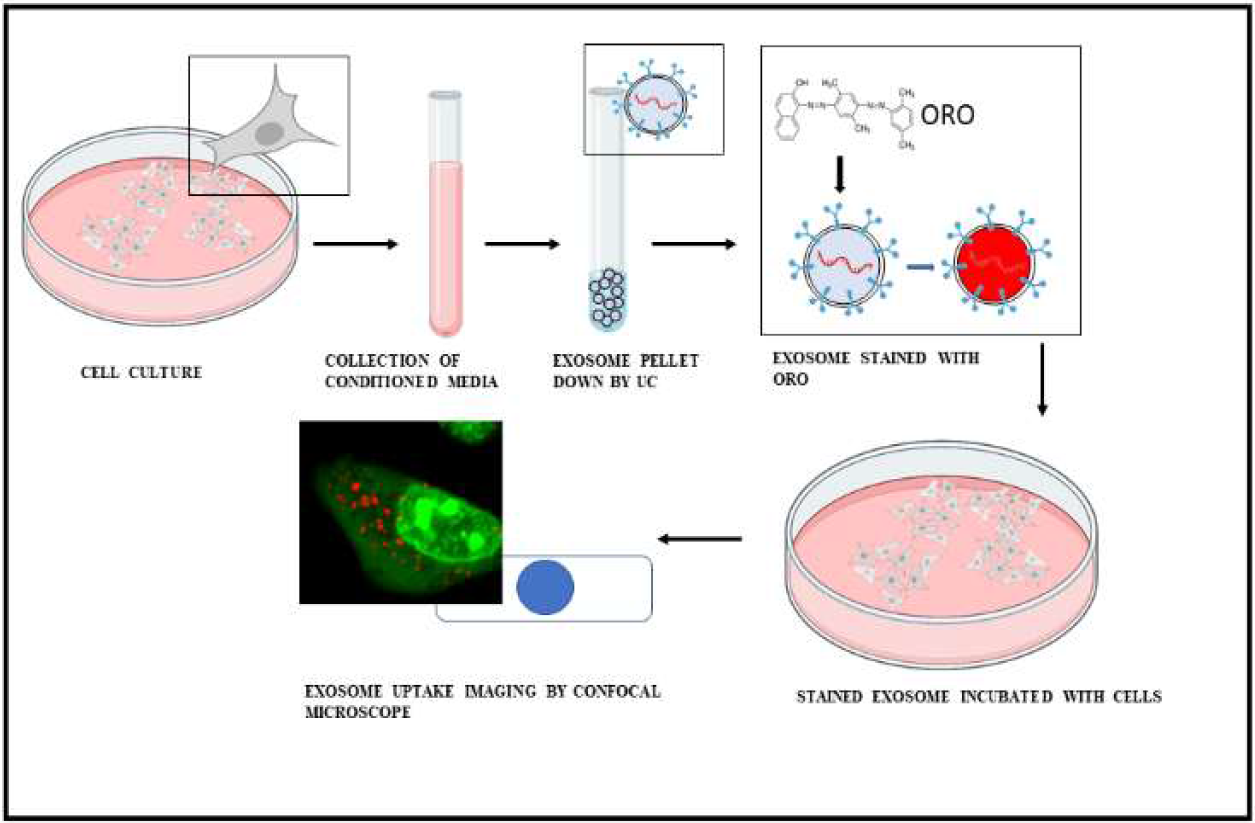

## Introduction

Exosomes are vesicles of 40–150 nm diameter released into the extracellular milieu by all cells. They are of endosomal origin formed by the fusion of multivesicular bodies with the plasma membrane. Structurally, exosomes are lipid bilayer vesicles containing nucleic acids, proteins amino acid, lipids and metabolites. They are stable entities with characteristic functions and activity [1, 2]. One of the important biological functions of exosomes is intercellular communication. Exosomes are now emerging as potential diagnostic and therapeutic tools for the treatment of various diseases, including cancer as they modulate various cellular functions such as growth, proliferation, tumorigenesis, tumour metastasis and therapy resistance. To understand their specific functional role in the cellular processes, their imaging and tracking studies are important. It is however difficult to characterize exosomes because of their small size and their complex nature. Their detection, purification and analysis are the major challenges for detailed study of their function and characterization.

Numerous Labelling and Detection methods have been used to examine exosome uptake into target cells and *in vivo* bio-distribution [3]. One of the widely used methods of exosomes labelling is the use of lipophilic fluorescent dyes that can integrate with the membrane of exosomes. These labelled exosomes can be detected using FLI in the visible light spectrum (390–700 nm). Commonly used organic fluorescent dyes originally used for labelling cell membrane are used for exosome labelling application. Examples of fluorescent lipid membrane (lipophilic) dyes widely used are PKH26, PKH67, DiI and DiD, and R18. The fluorophores of these dyes bind to different functional groups in the lipid bilayer to label the exosome membrane. Structurally, the PKH family of dyes has a polar head which is highly fluorescent and a long aliphatic tail which intercalates and binds to lipids. This leads to a strong fluorescence which remains stable and persists for a long duration [4]. DiI and DiR are a family of carbocyanine dyes that fluoresce strongly only when intercalated to a lipid-membrane. Structurally, they have two indoline rings with a long alkyl chain tail on each of the nitrogen and a conjugated 3-carbon bridge between the two aromatic rings. Because of their intercalation to the lipids, these dyes are fairly resistant to fading [5]. Octadecyl rhodamine B chloride (R18) is also a lipid dye that inserts into lipid membrane via its alkyl tails. R18 used in its quenched form to label membrane shows increased fluorescence when there is fusion between the labelled and unlabelled membrane. This has been used as a way of reporting exosomes fusion with cells [6].

The advantage of using the dyes mentioned above is low background signals since there is no transfer of dyes released from exosomes to unlabelled cells/tissues [7]. These dyes also give strong and stable signal for exosome detection. However, they have certain drawbacks such as long-term dye retention due to their intercalation with the lipid bilayer, formation of dye aggregates without exosomes, difficulty to distinguish dye particles from labelled exosomes and detection of large particles that are either aggregated dye or dye-exosome aggregates, which are taken up by cells with lesser efficiency than smaller exosomes.

The bio imaging modalities used for the detection of labelled exosomes are bioluminescence imaging (BLI), fluorescence imaging (FLI), nuclear imaging, and magnetic resonance imaging. BLI is a protein-based imaging where bioluminescent proteins, luciferases are used as reporters. These reporters oxidize their appropriate substrates to give bioluminescence. It has the highest sensitivity and high signal-to-noise ratio. However, BLI signal detection requires ultra-sensitive CCD camera, in addition to administration of substrates for luciferase. BLI also has low spatial and temporal resolution. FLI is relatively simpler with easy detection using a CCD camera and can be applied for real-time observation of exosomes. Fluorescence imaging involves the excitation of a fluorescent protein or organic dye with an external light source, which then emits signals that can be captured. It has high spatial resolution but low penetration [7]. Nuclear imaging shows the highest penetration with high sensitivity but it utilizes hazardous radioisotopes such as 99mTc and also has low spatial resolution and is expensive. Along with high spatial and temporal resolution, MRI has high penetration. However, MRI has low sensitivity and is expensive.

We report here for the first time ORO staining for exosomes. This is new, cost effective and easy approach to stain exosomes with intense red colouration. We evaluated ORO, a cell permeable dye, routinely used for intracellular lipid and tissue staining, for exosome labelling. A fat-soluble fluorescent dye, it stains neutral lipids, cholesterol esters and lipoproteins. It is an azo dye; and contains two azo groups attached to three aromatic rings. It is difficult to ionize, which renders it highly soluble in lipids. It stains lipids red and light absorption maximum is 518 nm. ORO stains neutral lipids (mainly triglycerides) with an orange-red tint. The principle of staining lipids with ORO is based on its higher solubility in lipids compared to the dye solvents. It does not form any bonds with the lipids and as such it is not truly a lipid stain. It simply moves from the dye solvent to the lipid due to its higher solubility in the latter. Further, the issues faced with commonly used lipophilic dyes for exosomes labelling, such as long term persistence of dyes, aggregation, micelle, difficulty to distinguish dye particles from labelled exosomes and detection of large aggregates of dye or dye-exosome are not seen with ORO staining. This method is advantageous over the existing methods as it shows good labelling efficiency with very sensitive detection and real-time tracking of the cellular uptake of exosomes. ORO also solves the issue of long -term persistence of commonly used lipophilic dyes.

## Result and Discussion

The exosomes were isolated from human A549 lung cancer cell lines grown in complete RPMI cell culture media. Cells growing in log phase were trypsinized and resuspended thoroughly to obtain a single-cell suspension. 2 × 10^5^ cells were seeded in 100 mm tissue culture petri-plates in triplicates in complete growth media. When cells were 40% confluent, media was removed, cells washed thrice with 1X PBS. Then exosomes depleted media was added to each plate. After 48 hours of incubation, media were collected and exosomes were isolated via ultracentrifugation method. The pellet containing exosomes was dissolved in 50 μl of 1XPBS.

The freshly isolated exosomes were characterized using TEM, AFM and DLS. To examine the morphology exosomes samples were spotted on a TEM copper grid for ∼2 mins. The samples were washed with water and then stained with 2% (w/v) aqueous uranyl acetate solution (*17*) for ∼10 min followed by another washing step. Air-dried grids were then examined at an accelerating voltage of 120 KV. Transmission electron microscopy (HR-TEM JEOL 2100F) images of the isolated exosomes showed spherical morphology [8]. The size of the exosomes was smaller than 100 nm (Figure 1a)]. Exosomes of similar sizes have been reported in other studies in A549 as well as other cancer cell lines [8 -11].

**Fig 1.**
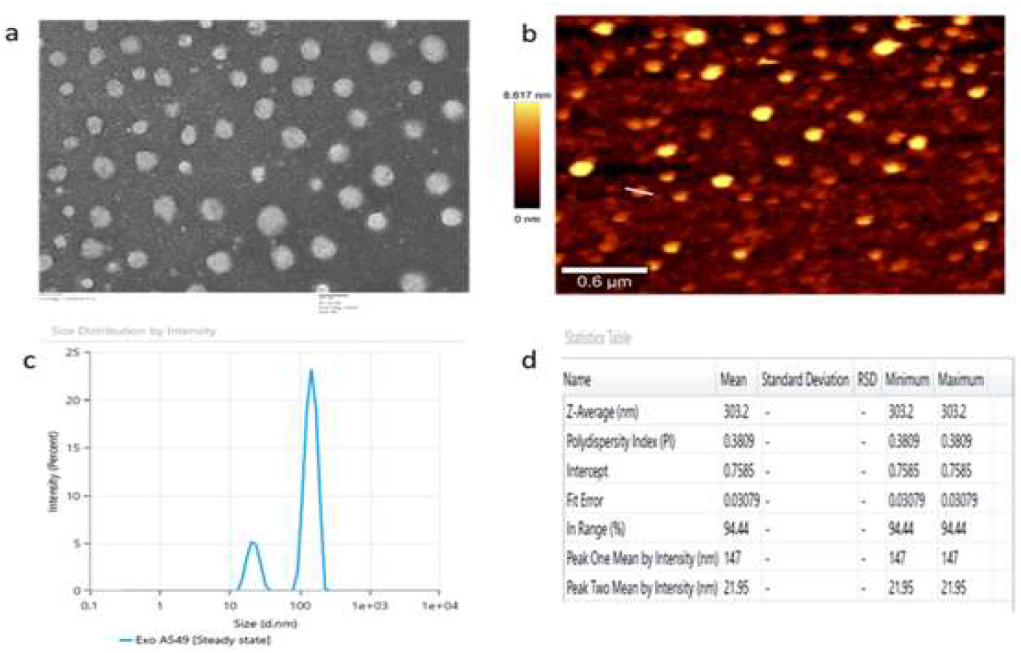
Characterization of isolated exosomes from A549 cells. (a) Transmission electron microscopy image of the isolated particles (b) Atomic force microscopy images showing topography [maximum height range of 8.617nm]. (c) &(d) DLS result for the size distribution of exosomes.

The surface morphology of the exosomes was visualized by conventional atomic force microscopy, using AFM system (WITec Germany). The samples were diluted (10 folds) in ultrapure water and 20μl samples smeared on freshly cleaved mica and air dried. The dried samples were then washed thoroughly with ultrapure water and air dried again. Images were taken immediately using tapping mode (NC-AFM) with a resonance frequency of 300 Hz. All AFM images were captured under ambient condition. Atomic Force Microscopy confirmed the spherical shape and the membrane integrity of the exosomes. The exosomes showed maximum height of 8.617 nm (Figure 1b). The size of the exosomes in terms of hydrodynamic size was measured using Dynamic Light Scattering. The DLS measurement was performed by spectrosize300 from Nano Bio chem Technology, Hamburg equipped with an inbuilt Peltier controller unit. Samples were resuspended in PBS before analysis. The Z average or overall mean size of the exosomes was found to be 303 nm (Figure 1c and 1d). The PDI was 0.3 which is within the acceptable range.

ORO stock-solution was prepared by dissolving 5 mg of dye in 1 ml of 100% isopropanol, and left overnight at room temperature. Next day, stain was filtered with 0.2 μm pore size membrane filter. A working solution of was made by mixing 350 μl ORO stock solution with 150 μl double distilled water. The working solution was again filtered with 0.1 μm membrane filter. The working concentration of ORO used to stain exosomes was 1.4μg/ml. Binding of ORO dye to exosomes was monitored by fluorescence spectrophotometer (Shimadzu RF-5300, Japan). The excitation and detection wavelengths were 560 nm and 690 nm respectively. The fluorescence intensity of ORO dye increased significantly after binding to exosomes. Fluorescence intensity was measured using arbitrary units. Increasing the amount of exosomes increased the fluorescence further (Fig. 2a).

**Fig 2.**
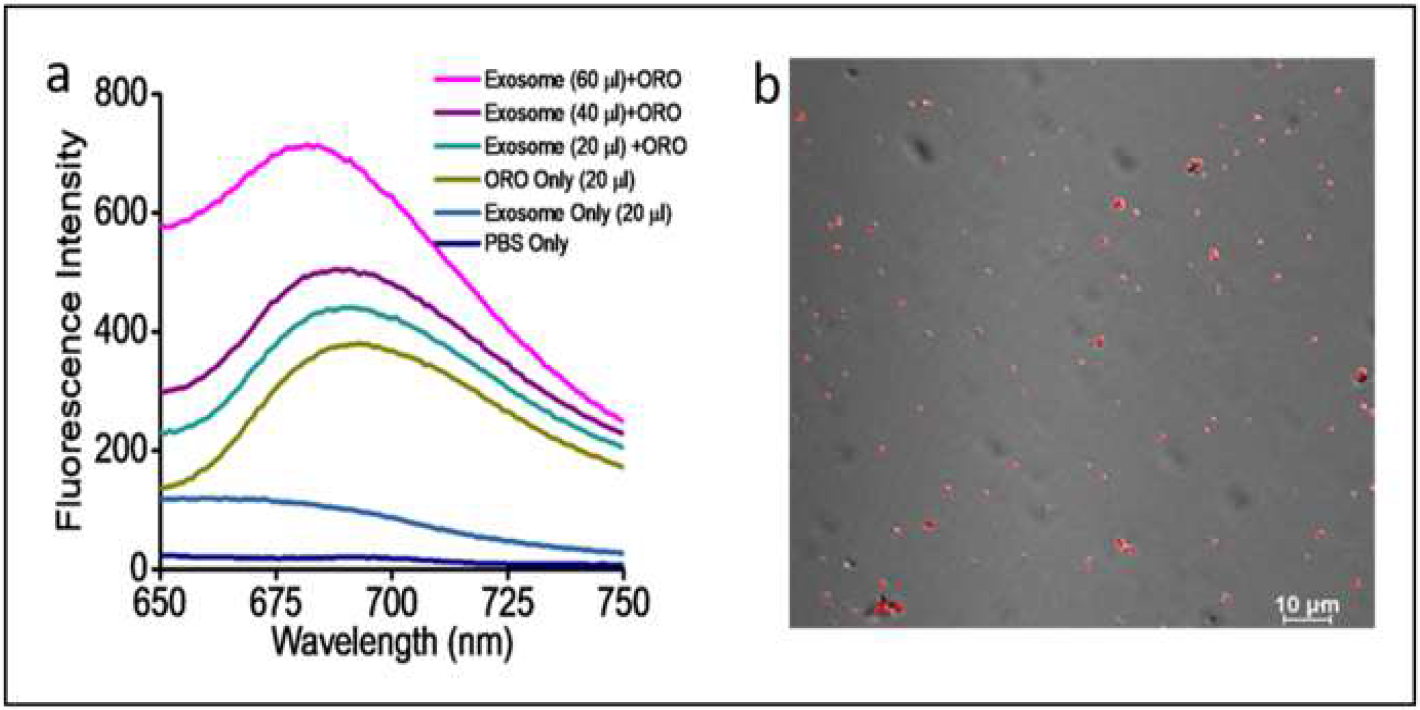
Staining of exosome by ORO a) Fluorescence emission spectra of Exosomes–Oil Red O complex. Fluorescence intensity increases with increase in exosome concentration (l_em_=690 nm and l_ex_=560 nm.). b) Isolated exosomes stained by ORO showing fluorescence under confocal microscopy.

ORO stained exosomes were also visualized under Confocal Microscope (Nikon AIR HD) Fig (2b). ORO may bind to exosomes cargo like lipids, proteins, RNA or even dsDNA to give red fluorescence [12].

To monitor uptake of labelled exosome by cells, ORO stained exosomes were incubated for 3 h with the cells grown on cover slips. Only exosomes stained with ORO showed fluorescence under TRIT C channel, while no fluorescence was detected in cells stained with same concentration of dye. (Supplementary fig. 1a and 1b). Incubation of cells with ORO stained exosomes upto 3h showed intracellular fluorescence. To further confirm that uptake of exosomes cells was stained with Acridine Orange for whole cell staining and Hoechst 33342 for nuclear staining. The exosomes were seen scattered in the cytoplasm and not inside the nucleus (Fig. 3a and 3b).

**Fig 3.**
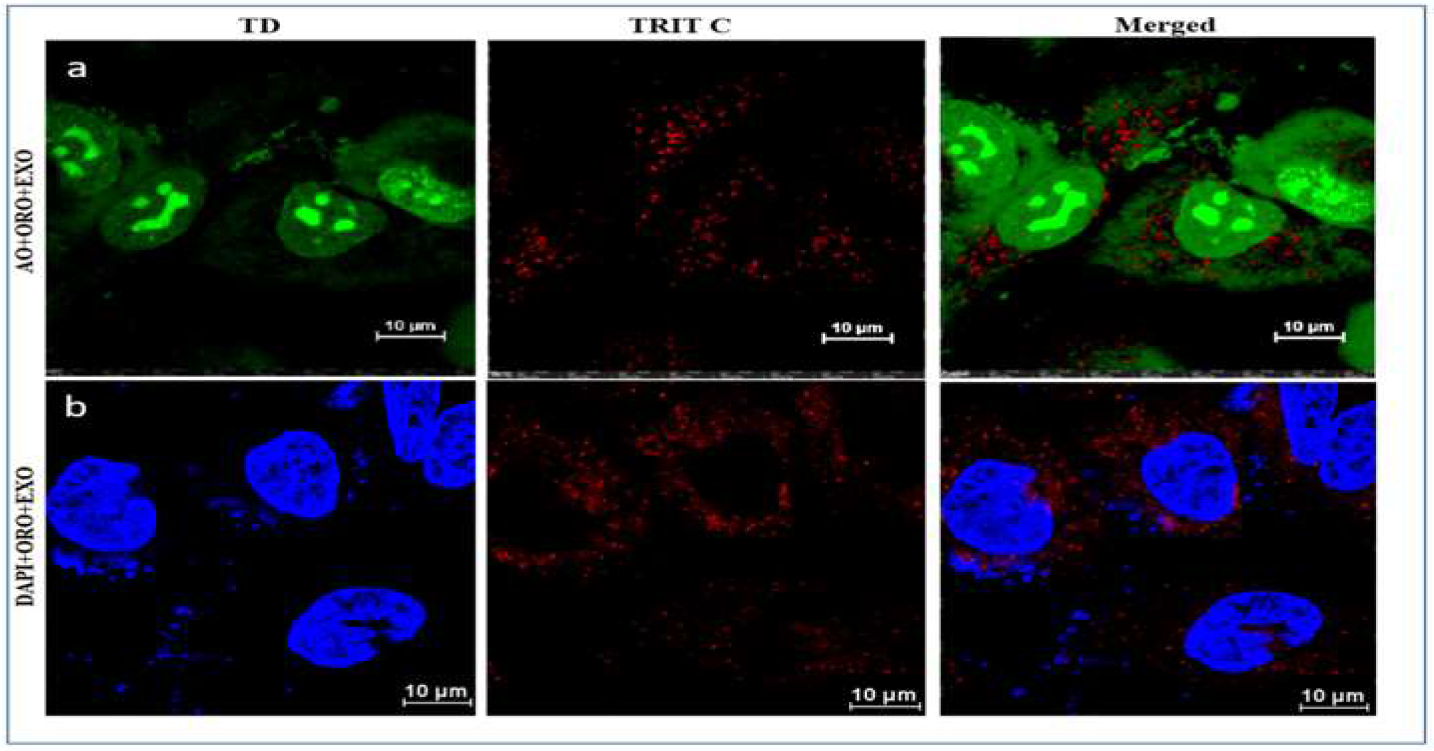
Exosome localization and uptake by A549 cells incubated with ORO stained exosomes. a)Cells stained green (acridine orange). b) Nucleus stained blue (Hoechest). All confocal microscopy images were captured under 100 X (Scale: 10 μm).

To further check the binding of dye to cells, cell was incubated with same concentration of dye and incubated for the same time points without exosomes. Cells treated with this concentration of ORO did not stain any cell components (Supplementary fig. 2a and 2b). The intracellular cytoplasmic localization of ORO stained exosomes was further confirmed by the 3D Confocal Microscopy Z slicing images (Figure 4). This indicates the effective localization of exosomes inside the cell.

**Fig 4.**
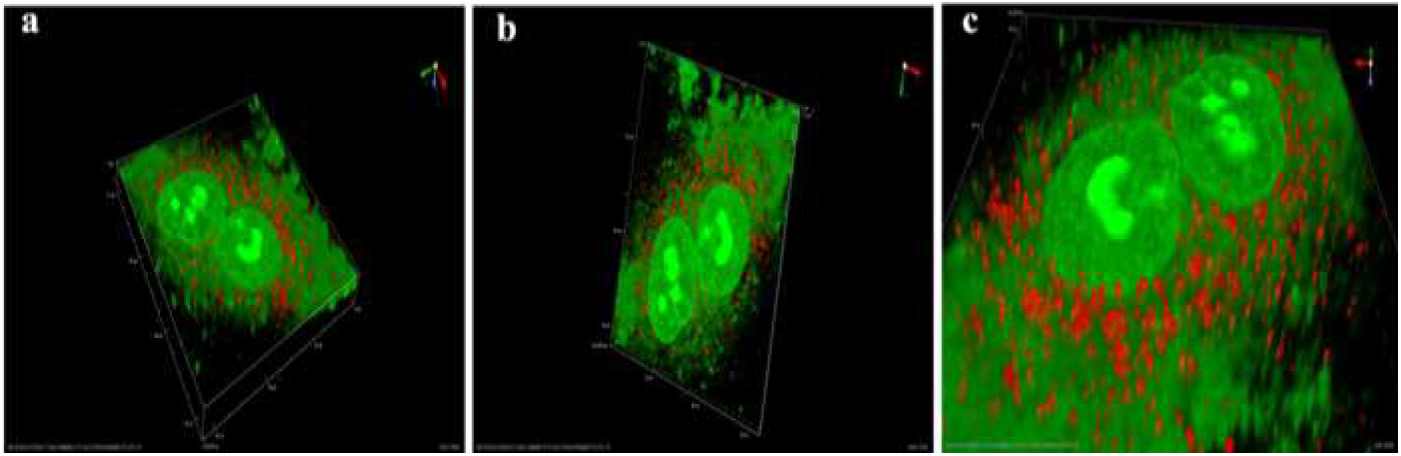
Representative Z-stack confocal image through the volume of the treated cells that demonstrate the internalization and localization of ORO (oil red o) stained exosomes in A549 cells. A volume rendered 3D construction of the fluorescent z-stack image (a) and (b) in different orientations, which further demonstrates the localization of exosomes in cell and cytoplasm. (c) Zoomed view of 3D reconstructed fluorescence image.

In conclusion, staining of exosome using ORO was done successfully, with good sensitivity of detection using fluorescence microscopy. This method did not show any dye aggregates; dye-exosome aggregates were also negligible. The dye binding to exosomes increased the fluorescence significantly, because of which it was easy to detect and track the exosome uptake by the cells. The concentration of dye required to stain exosomes does not stain any cell components making it easier to localize them in live cells. Uptake of the stained exosomes by live cells was easy to monitor because of the distinct red fluorescence. Thus, ORO staining method can be used as a good alternative to commonly used fluorescent dye for exosome labelling and uptake studies.

There are no conflicts to declare.

