## Supplementary figures and images for "Oil Red O based method for Exosome Labelling and detection"

### Supplementary fig 1.

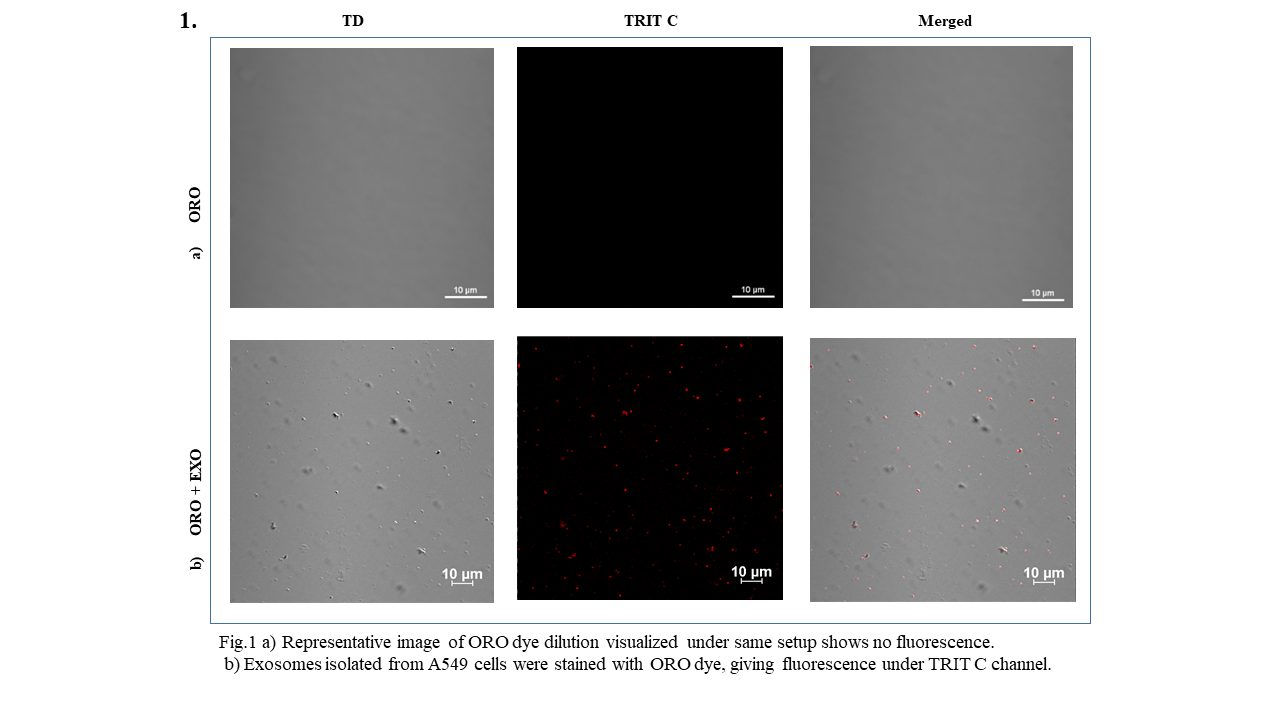

### Supplementary fig.2

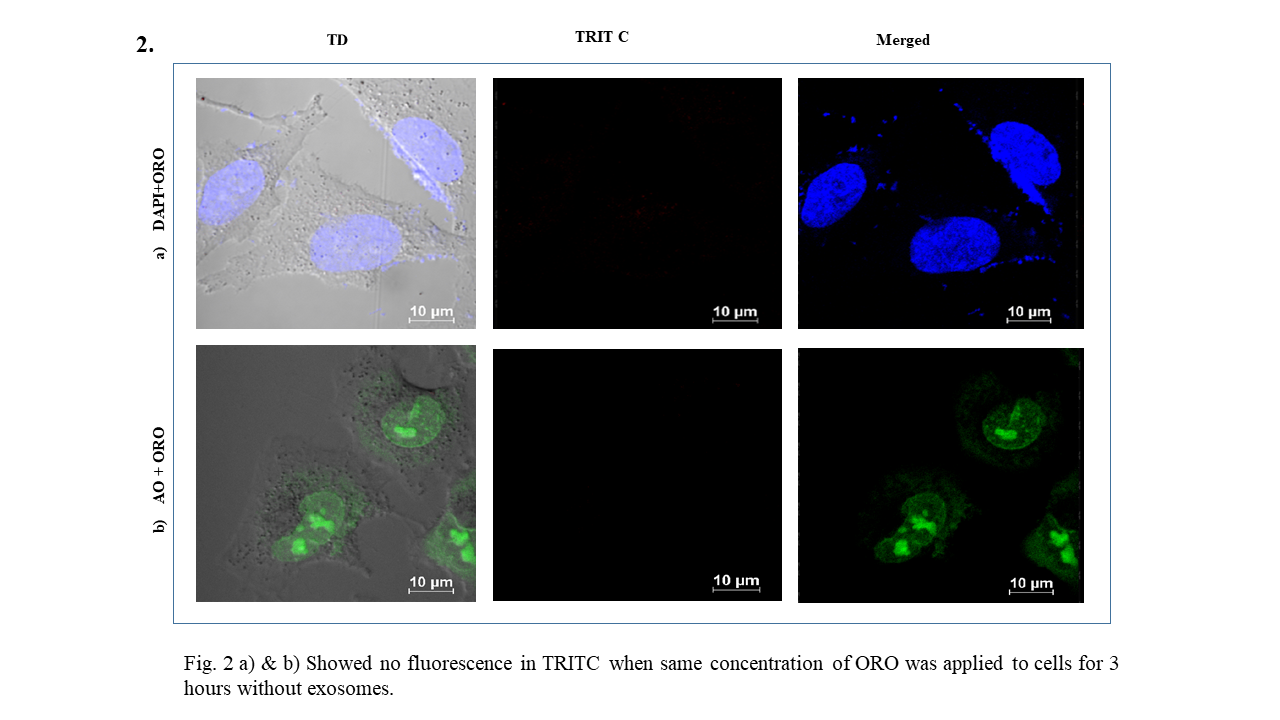
